# The natural stilbenoid (–)-hopeaphenol inhibits cellular entry of SARS-CoV-2 USA-WA1/2020, B.1.1.7 and B.1.351 variants

**DOI:** 10.1101/2021.04.29.442010

**Authors:** Ian Tietjen, Joel Cassel, Emery T. Register, Xiang Yang Zhou, Troy E. Messick, Frederick Keeney, Lily D. Lu, Karren D. Beattie, Topul Rali, Hildegund C. J. Ertl, Joseph M. Salvino, Rohan A. Davis, Luis J. Montaner

## Abstract

Antivirals are urgently needed to combat the global SARS-CoV-2/COVID-19 pandemic, supplement existing vaccine efforts, and target emerging SARS-CoV-2 variants of concern. Small molecules that interfere with binding of the viral spike receptor binding domain (RBD) to the host ACE2 receptor may be effective inhibitors of SARS-CoV-2 cell entry. Here we screened 512 pure compounds derived from natural products using a high-throughput RBD/ACE2 binding assay and identified (–)-hopeaphenol, a resveratrol tetramer, in addition to vatalbinoside A and vaticanol B, as potent and selective inhibitors of RBD/ACE2 binding and viral entry. For example, (–)-hopeaphenol disrupted RBD/ACE2 binding with a 50% inhibitory concentration (IC50) of 0.11 μM in contrast to an IC50 of 28.3 μM against the unrelated host ligand/receptor binding pair PD-1/PD-L1 (selectivity index = 257.3). When assessed against the USA-WA1/2020 variant, (–)-hopeaphenol also inhibited entry of a VSVΔG-GFP reporter pseudovirus expressing SARS-CoV-2 spike into ACE2-expressing Vero-E6 cells and *in vitro* replication of infectious virus in cytopathic effect assays (IC50 = 10.2 μM) without cytotoxicity. Notably, (–)- hopeaphenol also inhibited two emerging variants of concern originating from the United Kingdom (B.1.1.7) and South Africa (B.1.351) in both cytopathic effect and spike-containing pseudovirus assays with similar (B.1.1.7) or improved (B.1.351) efficacies over the USA- WA1/2020 variant. These results identify (–)-hopeaphenol and related stilbenoid analogues as potent and selective inhibitors of viral entry across multiple SARS-CoV-2 variants including those with increased infectivity and/or reduced susceptibility to existing vaccines.

**Importance:** SARS-CoV-2 antivirals are needed to supplement existing vaccine efforts and target emerging viral variants with increased infectivity or reduced susceptibility to existing vaccines. Here we show that (–)-hopeaphenol, a naturally-occurring stilbenoid compound, in addition to its analogues vatalbinoside A and vaticanol B, inhibit SARS-CoV-2 by blocking the interaction of the viral spike protein with the cellular ACE2 entry receptor. Importantly, in addition to inhibiting the early USA-WA1/2020 SARS-CoV-2 variant, hopeaphenol also inhibits emerging variants of concern including B.1.1.7 (“United Kingdom variant”) and B.1.351 (“South Africa variant”), with improved efficacy against B.1.351. (–)-Hopeaphenol therefore represents a new antiviral lead against infection from multiple SARS-CoV-2 variants.

## Introduction

Severe acute respiratory syndrome coronavirus 2 (SARS-CoV-2) is the causative agent of Coronavirus Disease 2019 (COVID-19). Since crossing into humans in late 2019, SARS-CoV-2 has continued to cause substantial human morbidity and mortality worldwide. While SARS-CoV-2 vaccines are in development with several approved for emergency use, access to these vaccines remains limited, particularly in low and middle-income countries. Moreover, vaccine hesitancy and ongoing mutation of SARS-CoV-2 increase the risk of vaccine resistance. Finally, no reliable therapeutics are currently available to combat SARS-CoV-2 in those that have already developed COVID-19 or to mitigate SARS-CoV-2 spread to exposed individuals. Thus, SARS-CoV-2 antivirals are urgently needed to complement ongoing vaccination efforts.

One attractive therapeutic target of SARS-CoV-2 replication is its binding and entry into host cells, which is induced by the trimeric viral spike glycoprotein (1). A primary cellular receptor of SARS-CoV-2 entry is the angiotensin-converting enzyme II (ACE2) protein. Viral entry is mediated by the receptor biding domain (RBD) of the S1 segment of spike, which directly interacts with ACE2, while S2 mediates membrane fusion (2–3). Following RBD/ACE2 binding, SARS-CoV-2 gains entry to host cells through both an endosomal, clathrin-dependent pathway as well as a clathrin-independent pathway which involves spike protein cleavage by furin, TMPRSS2, and other host proteases (1–2). Antagonism of the RBD/ACE2 interaction would therefore be expected to block SARS-CoV-2 entry and replication.

Since its worldwide outbreak in early 2020, SARS-CoV-2 variants have acquired mutations within Spike that enhance binding to ACE2 and increase infectivity, as first detected by the emergence of a D614G mutation which has since spread worldwide (4). More recently, additional variants have emerged with further spike sequence divergence; these include B.1.1.7, which originated in the United Kingdom, and B.1.351 (also called 501Y.V2) from South Africa (5). The B.1.1.7 variant contains an additional N501Y mutation within the RBD that causes reduced antibody neutralization by both convalescent and vaccine sera *in vitro* (6). Concerningly, the B.1.351 variant, which along with N501Y has also acquired mutations E484K and K417N within the RBD, has further reduced susceptibility to, or escape from, neutralizing antibodies as well as sera from convalescent and vaccine-treated patients and immunized mice (7–8). As these observations may raise future concerns about the long-term efficacy of existing vaccines, additional countermeasures with the potential to target emerging SARS-CoV-2 variants are essential.

Pure compounds derived from natural products are a rich source of antivirals including against coronaviruses (9). However, outside of computational studies, few natural products to date have been demonstrated to act on SARS-CoV-2 replication (10). To identify natural product-derived compounds that may inhibit entry across multiple SARS-CoV-2 variants, we developed an AlphaScreen-based RBD/ACE2 interaction assay to screen a pure compound library containing 512 natural products and derivatives sourced predominantly from plants, mushrooms and marine invertebrates of Australia, Papua New Guinea, and neighboring regions (11–12). The top hits from this screen, which included the plant stilbenoids (–)-hopeaphenol, vatalbinoside A, and vaticanol B (**Figure 1**), were then assessed for *in vitro* mechanisms of action using SARS-CoV-2 pseudoviruses and antiviral efficacy using infectious SARS-CoV-2 variants encompassing parental and B.1.1.7 and B.1.351 variants.

**Figure 1.**
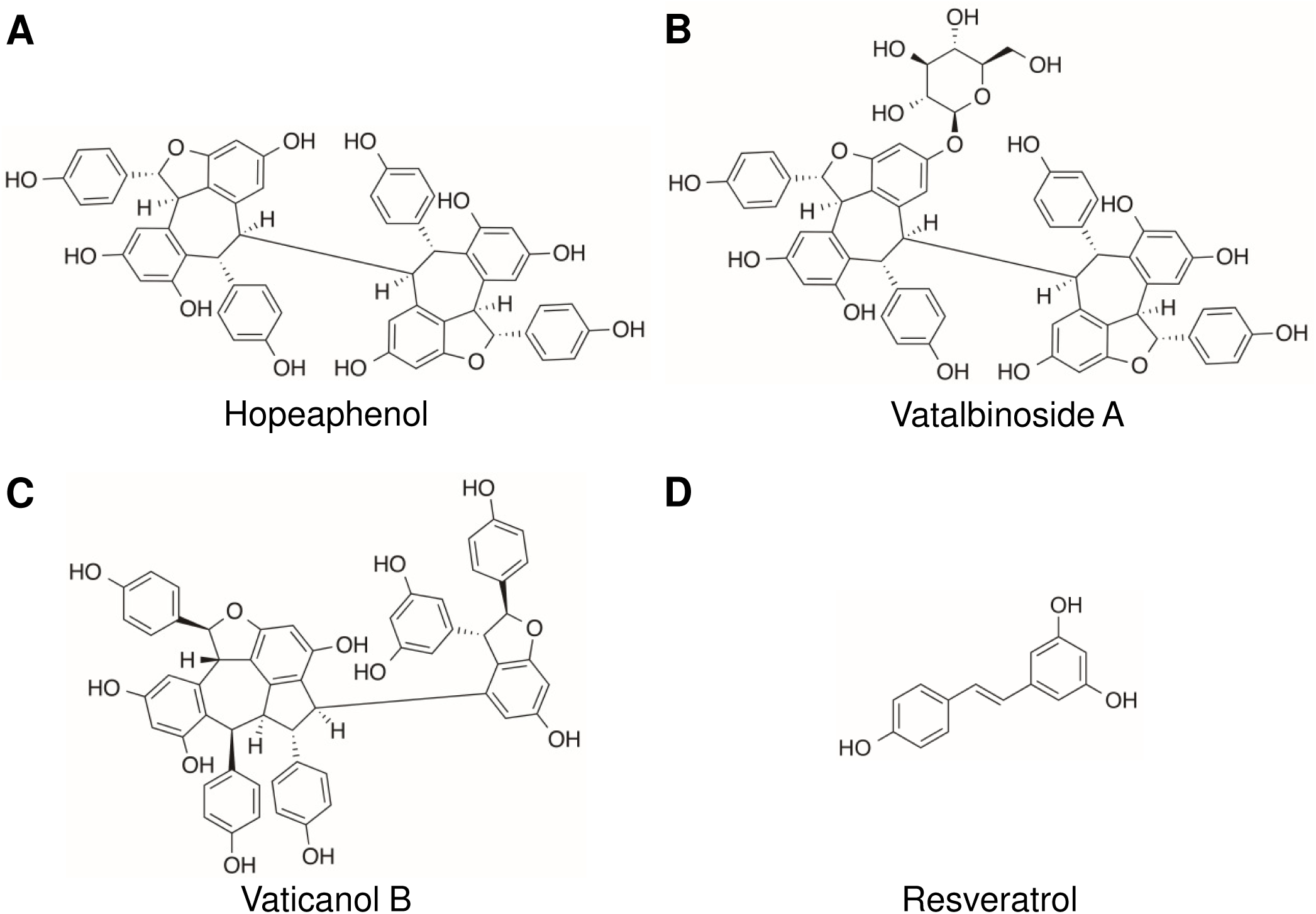
Chemical structures of (–)-hopeaphenol **(A)**, vatalbinoside A **(B)**, vaticanol B **(C)**, and resveratrol **(D)**.

## Results

### Stilbenoids selectively inhibit the SARS-CoV-2 spike RBD / host ACE2 protein interaction

To identify potential inhibitors of SARS-CoV-2 entry, we used AlphaScreen technology (13) to develop a high-throughput, 384 well plate-based assay to monitor the interaction of SARS-CoV-2 Spike RBD with host ACE2 (**Figure 2A**). Briefly, a SARS-CoV-2 RBD protein derived from USA-WA1/2020 and containing a C-terminal His tag, in addition to a full-length ACE2 peptide with a C-terminal Fc tag, were pre-bound to respective acceptor and donor beads and co-incubated for 3 hours at room temperature. When a ligand/receptor binding event occurs, excitation at 680 nm results in a singlet oxygen transfer between donor and receptor beads, which results in luminescence at 615 nm. Compounds that inhibit binding of RBD to ACE2 should therefore inhibit luminescence. In the absence of compounds, we observed that luminescence was highly dependent on concentrations of both RBD and ACE2 (**Figure 2A**) with a > 200-fold signal to noise ratio and high reproducibility across experiments (Z’ > 0.8) (14).

**Figure 2.**
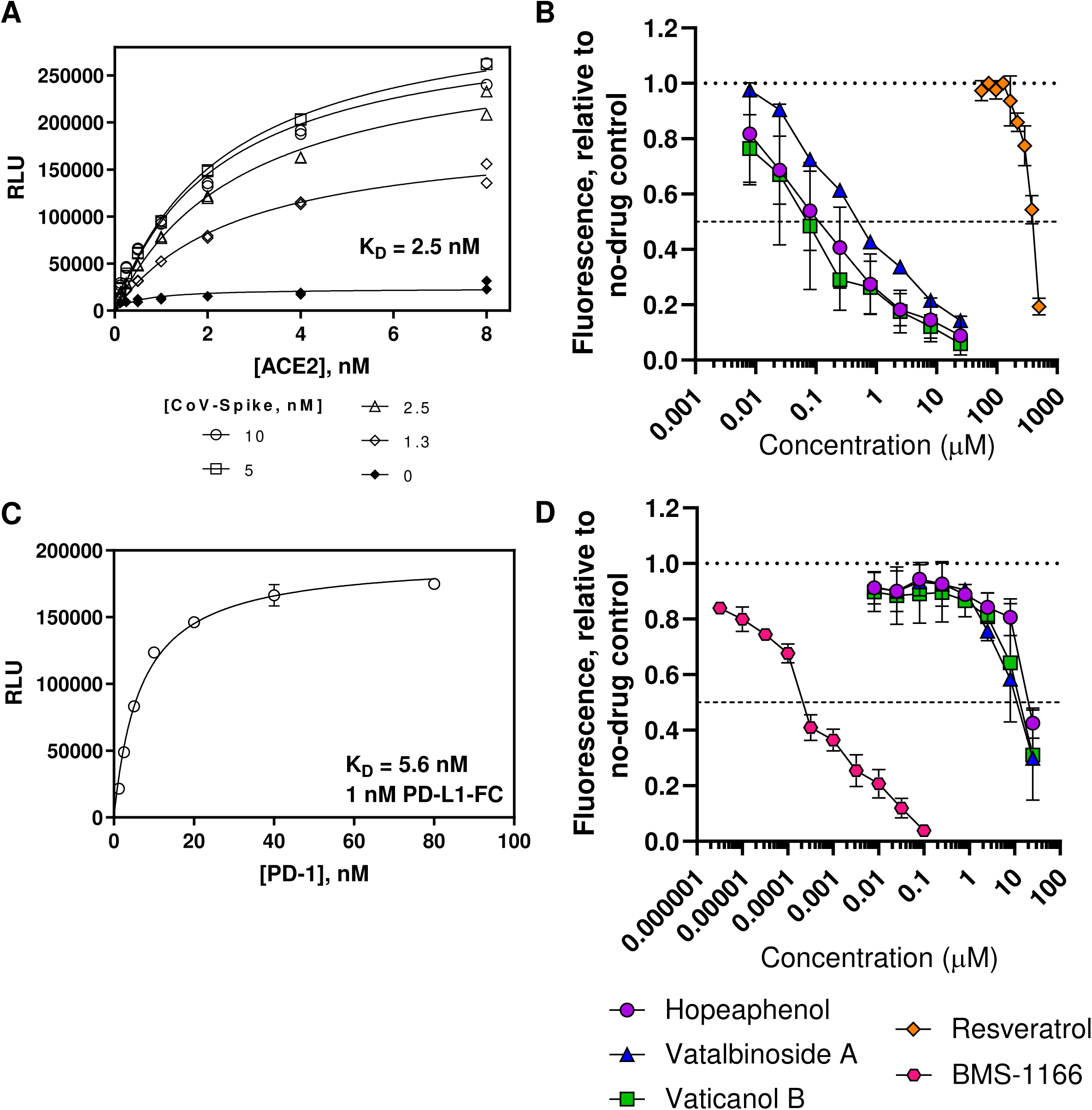
Identification of stilbenoids as Spike-ACE2 inhibitors by AlphaScreen. **A,** Demonstration of AlphaScreen-based fluorescence due to interactions of His-tagged Spike RBD (from USA-WA1/2020) and Fc-tagged ACE2 peptides. **B,** Dose-response curves of stilbenoids and resveratrol on fluorescence inhibition due to disruption of RBD/ACE2 interactions. **C,** Demonstration of AlphaScreen-based fluorescence due to interactions of His-tagged PD-1 and Fc-tagged PD-L1 peptides. **D,** Dose-response curves of stilbenoids and control inhibitor BMS-1166 on fluorescence inhibition due to disruption of PD-1/PD-L1 interactions.

Using this assay, we then screened 512 pure compounds obtained from natural products and semisynthetic derivatives available from Compounds Australia at Griffith University at 1 μM, where 9 (1.8%) inhibited > 75% of fluorescence observed in the absence of compounds. Activities of top compounds were then assessed for dose-response profiles, where the three most active hits were a series of stilbenoid resveratrol tetramers including (–)-hopeaphenol, vatalbinoside A, and vaticanol B (**Figure 1; Figure 2B**). These stilbenoids, exemplified by hopeaphenol, are tetramers of resveratrol available from multiple plant sources (15). Using the RBD/ACE2 AlphaScreen assay described above, dose-response profiles from 5 independent experiments were obtained to calculate 50% inhibitory concentrations (IC50s) of 0.11, 0.24, and 0.067 μM for hopeaphenol, vatalbinoside A, and vaticanol B, respectively (**Table 1**). In contrast, almost no activity was observed in this assay by the resveratrol monomer (IC50 > 100 μM; **Figure 2B**), indicating that inhibition requires a multimeric structure.

**Table 1.**
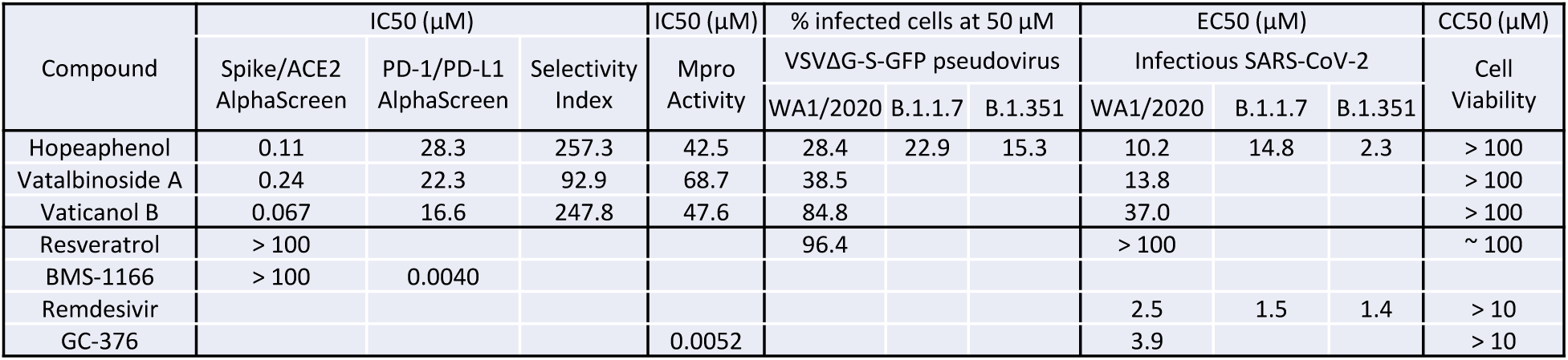
Summary of total stilbenoid and control compound bioactivities.

To confirm selectivity of hopeaphenol and analogues to disrupt the RBD/ACE2 interaction, we next assessed their ability to interfere with the unrelated host PD-1/PD-L1 ligand/receptor pair using a comparable and previously-described experimental approach (16), where we observed a > 100-fold signal to noise ratio and Z’ > 0.75 (**Figure 2C**). In this assay, the control PD-1/PD-L1 antagonist BMS-1166 (17) disrupted bead proximity-based fluorescence with an IC50 of 0.0040 μM (**Figure 2D**) but had no activity against the RBD/ACE2 interaction (IC50 > 100 μM; data not shown). Conversely, hopeaphenol, vatalbinoside A, and vaticinol B were all substantially less effective in disrupting PD-1/PD-L1, with IC50s of 28.3, 23.3, and 16.6 μM, respectively (**Figures 2D**; **Table 1**). From these two assays, the selectivity indices of hopeaphenol, vatalbinoside A, and vaticanol B [i.e., IC50 (PD-1/PD-L1) / IC50 (RBD/ACE2)] were calculated to be 257.3, 92.9, and 247.8, respectively, indicating high selectivity of these compounds for disrupting the viral RBD/host ACE2 interaction over an unrelated host ligand/receptor pair.

### Stilbenoids are weak inhibitors of viral main protease

A recent high-throughput virtual screening study proposed that hopeaphenol may act as an inhibitor of the SARS-CoV-2 main protease (M^pro^) by interfering with its active site (18), thereby raising the possibility of hopeaphenol acting on multiple viral targets. To test this possibility, we also developed an M^pro^ enzymatic assay using an M^pro^ peptide substrate resembling those described previously (19), where a C-terminal 5-((2-aminoethyl)amino)naphthalele-1-sulfonic acid (EDANS) fluorescent tag is quenched by an N-terminal 4-((4-(dimethylamino)phenyl)azo)benzoic acid (DABCYL) tag. Following incubation with recombinant M^pro^, the cleaved substrate affords separation of the EDANS tag from the DABCYL quencher and detection of fluorescence at 490 nm. Compounds that inhibit M^pro^ activity are therefore expected to inhibit fluorescence. This assay also exhibited a > 10-fold signal to noise ratio and Z’ > 0.6 (**Figure 3A**) and was adaptable to 384-well screening format.

**Figure 3.**
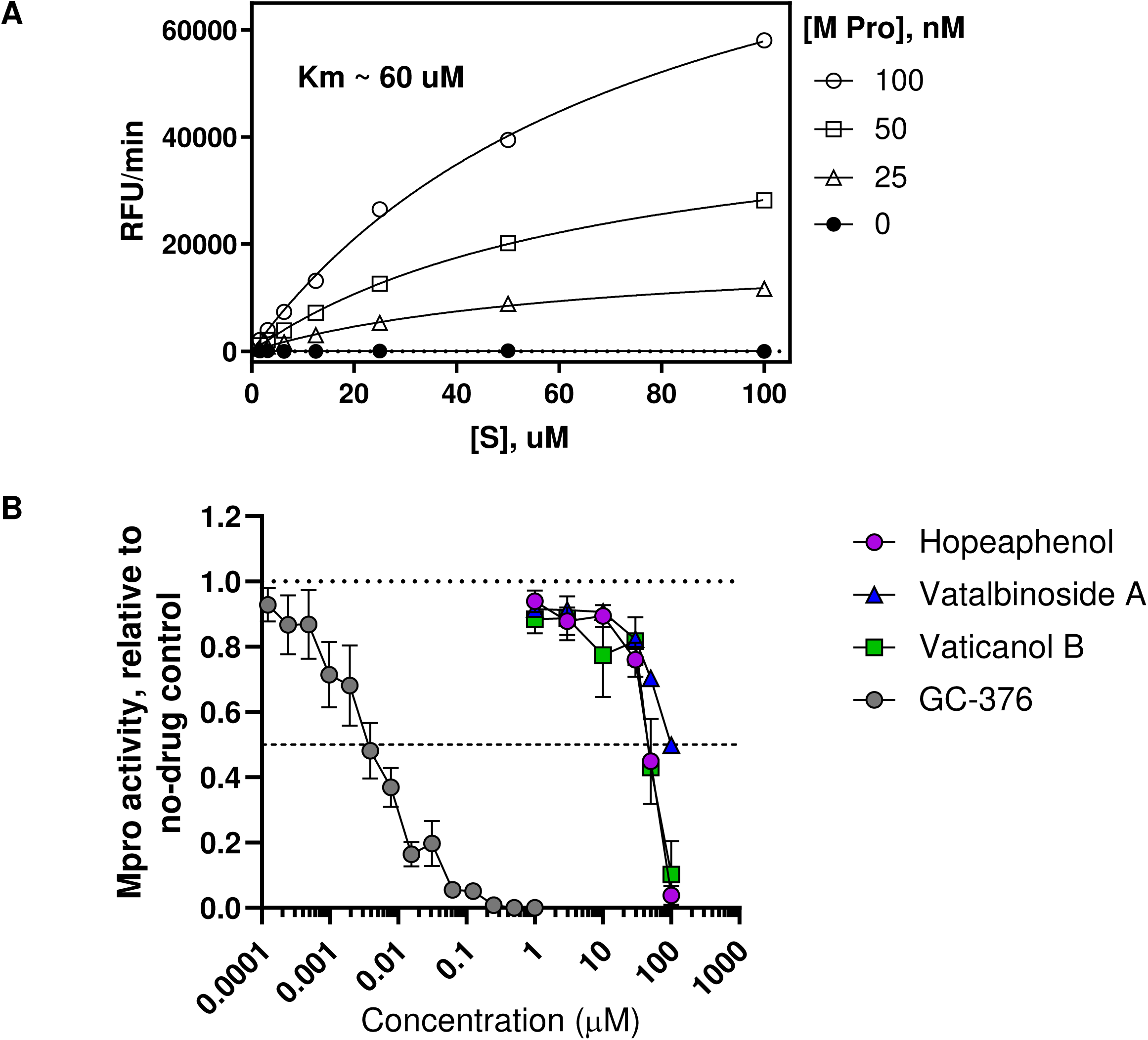
Effects of stilbenoids on inhibition of SARS-CoV-2 M^pro^ activity. **A,** Demonstration of recombinant M^pro^ enzymatic activity on a FRET-based fluorogenic peptide substrate. **B,** Dose-response curves of stilbenoids and control inhibitor GC-376 on M^pro^ enzymatic activity.

Using this assay, we observed that the control M^pro^ inhibitor GC-376 blocked enzymatic activity with an IC50 of 0.0052 μM, consistent with previous observations (**Figure 3B**) (19–20). In contrast, we observed that hopeaphenol, vatalbinoside A, and vaticanol B inhibited M^pro^ activity with IC50s of 42.5, 68.7 and 47.6 μM, respectively (**Figure 3B**; **Table 1**), suggesting that these stilbenoids, while potentially capable of targeting M^pro^, are to a first approximation more effective against RBD binding to ACE2.

### Stilbenoids inhibit SARS-CoV-2 spike-dependent viral entry

To assess whether hopeaphenol and analogues inhibit viral entry within a cellular context, we generated a single-cycle pseudovirus consisting of a vesicular stomatitis virus (VSV) backbone lacking the G fusion protein and expressing SARS-CoV-2 spike protein and green fluorescent protein (GFP) reporter (VSVΔG-S-GFP) (21). In our initial experiments, we generated pseudovirus with spike from the USA-WA1/2020 variant. Pseudovirus was then incubated with cells in the presence or absence of stilbenoids. High-content imaging was then used to count total live and infected cells in each culture, as determined by Hoechst-stained nuclei and cellular GFP fluorescence, respectively. Consistent with previous observations (22–23), VSVΔG-S-GFP pseudovirus infected ACE2-expressing cells like Vero-E6 (**Figure 4A**) (24) but not cell lines lacking ACE2 like BHK-21 cells (data not shown). In this assay, we observed an average of 2.5 ± 0.1% GFP-positive cells (mean ± s.e.m.) following 24 hours’ incubation with pseudovirus and 0.1% DMSO vehicle control (**Figure 4A**) with no major changes in number of cell nuclei relative to uninfected cells (**Figure 4B**). Additionally, no major changes in total cell nuclei were observed in the presence of up to 50 μM of any compound, indicating no overt effects on cell viability, with the exception of 50 μM resveratrol which resulted in cultures with 60.3 ± 7.0% of nuclei observed in untreated, infected cells (**Figure 4B**).

**Figure 4.**
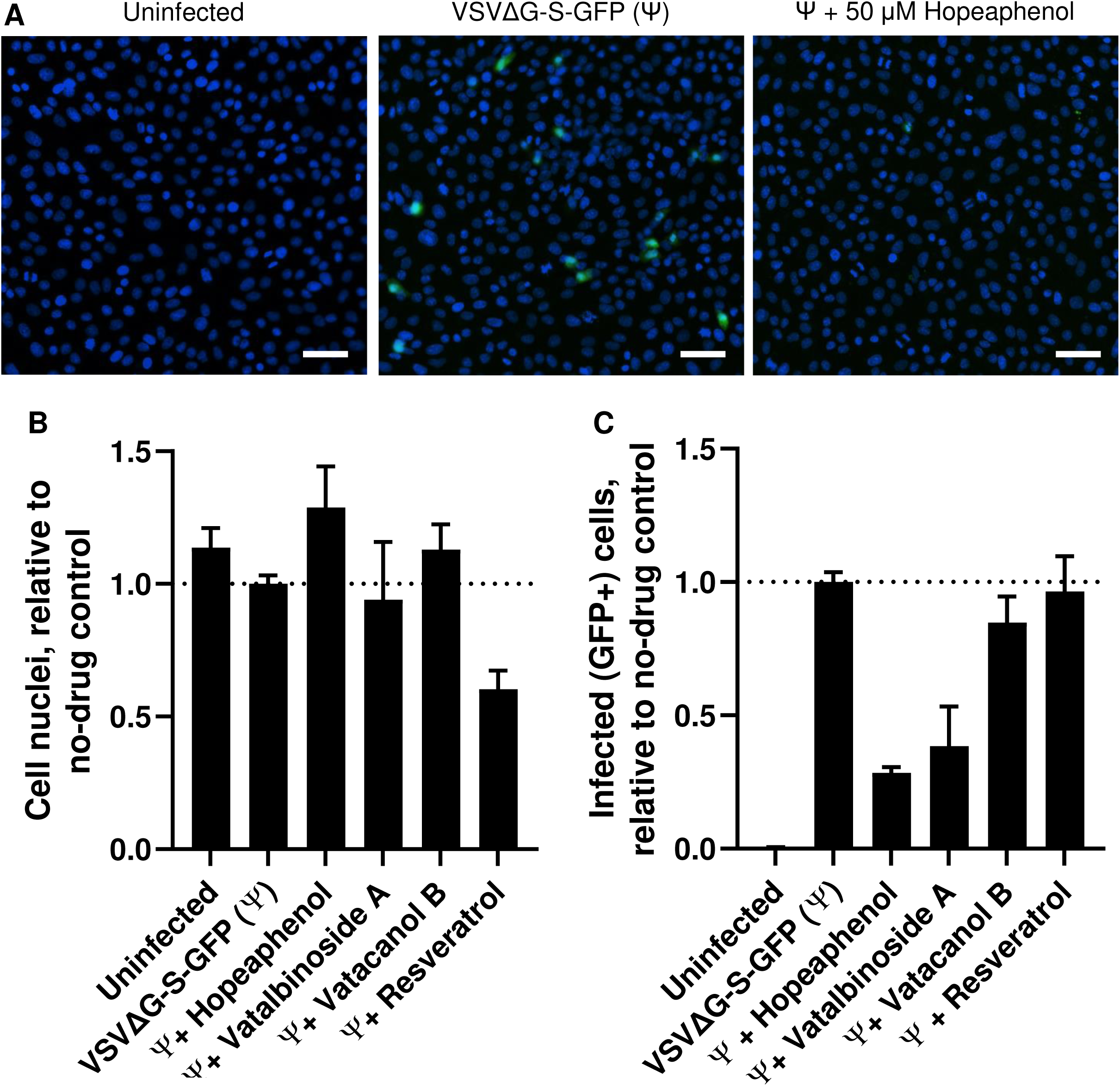
Effects of stilbenoids on *in vitro* pseudovirus entry. **A,** Representative images of uninfected Vero-E6 cells (left) or cells infected with VSVΔG-S-GFP pseudovirus containing SARS-CoV-2 Spike (USA-WA1/2020) in the absence (center) or presence of 50 μM hopeaphenol (right). Blue denotes Hoechst-stained cell nuclei, and green indicates GFP-positive infected cells. Scale bar = 100 μm. **B,** Average number of total cell nuclei per cell field as counted by high-content imaging. **C,** Level of GFP-positive (i.e., pseudovirus-infected) cells as measured by high-content imaging, relative to total number of cell nuclei, in the presence of stilbenoids. In **B** and **C**, data are presented relative to pseudovirus-infected cells in the presence of 0.1% DMSO vehicle control.

However, when Vero-E6 cells were infected with pseudovirus in the presence of 50 μM hopeaphenol, GFP fluorescence was present in only 28.4 ± 2.1% of infected cells treated with 0.1% DMSO (**Figure 4A, C**). Similar results were observed when infected cells were co-treated with 50 μM vatalbinoside A, where 38.5 ± 14.8% of GFP-positive cells were observed relative to infected, vehicle-treated cells (**Figure 4C**). In contrast, 50 μM vaticanol B resulted in GFP expression in 84.8 ± 9.7% of cells, indicating that the potent anti-RBD/ACE2 activity observed by AlphaScreen assay was not reproduced in the pseudotype assay. As expected, 50 μM resveratrol had no effect on GFP-positive cells (96.4 ± 13.3% GFP expression of infected, vehicle-treated cells; **Figure 4C**). However, no compound inhibited GFP expression when incubated with infected cells at 15 μM (data not shown). Taken together, these results indicate that at least a subset of stilbenoids can inhibit entry of pseudoviruses expressing SARS-CoV-2 spike protein *in vitro*, consistent with AlphaScreen assay results, although this occurs at much higher concentrations.

### Stilbenoids inhibit infectious SARS-CoV-2 replication

To confirm cellular antiviral activity of hopeaphenol and analogues, we next used a cytopathic effect (CPE)-based assay with infectious virus in Vero-E6 cells (25–26). Briefly, Vero-E6 cells were treated with compounds for 2 hours in 8-fold replicates in 96-well format before infection with 50x median tissue culture infectious dose (TCID50) of SARS-CoV-2 (USA-WA1/2020 variant). Cells were then incubated for 4 days with daily scoring of CPE across all wells by a user blinded to experimental conditions. Using this approach, we observed the presence of CPE by 2 days post infection, as characterized by extensive cell rounding and cellular debris that were observable by light microscopy (**Figure 5A**, arrows). By 4 days post-infection, this CPE was widespread across the cell culture and clearly distinguishable from uninfected cell controls (**Figure 5A**). When low micromolar concentrations of either the control nucleoside analog remdesivir or the M^pro^ inhibitor GC-376 (27–28) were added 2 hours before infection, CPE was completely inhibited in these cultures after 4 days (**Figure 5B**, top and middle), with calculated EC50s of 2.5 and 3.9 μM for remdesivir and GC-376, respectively (**Figure 5C**; **Table 1**). Moreover, comparable activity was observed in the presence of hopeaphenol (**Figure 5B**, bottom), which blocked SARS-CoV-2 replication after 4 days with a calculated EC50 of 10.2 μM (**Figure 5C**; **Table 1**). Notably, while similar antiviral activity was observed with vatalbinoside A (EC50 = 13.8 μM), we observed substantially less efficacy by vaticanol B (EC50 = 37.0 μM; **Figure 5C**; **Table 1**), consistent with its reduced efficacy in pseudovirus assays (**Figure 4C**). In contrast, no antiviral activity was observed by up to 100 μM resveratrol (**Figure 5C**). We also observed no evidence of cytotoxicity by these compounds, as measured by resazurin staining following 4 days treatment of uninfected Vero-E6 cells (**Figure 5D**). These results indicate that hopeaphenol and vatalbinoside A, and to a lesser extent vaticanol B, inhibit SARS-CoV-2 replication *in vitro*, with efficacy of hopeaphenol at the same order of magnitude as control SARS-CoV-2 antivirals remdesivir and GC-376.

**Figure 5.**
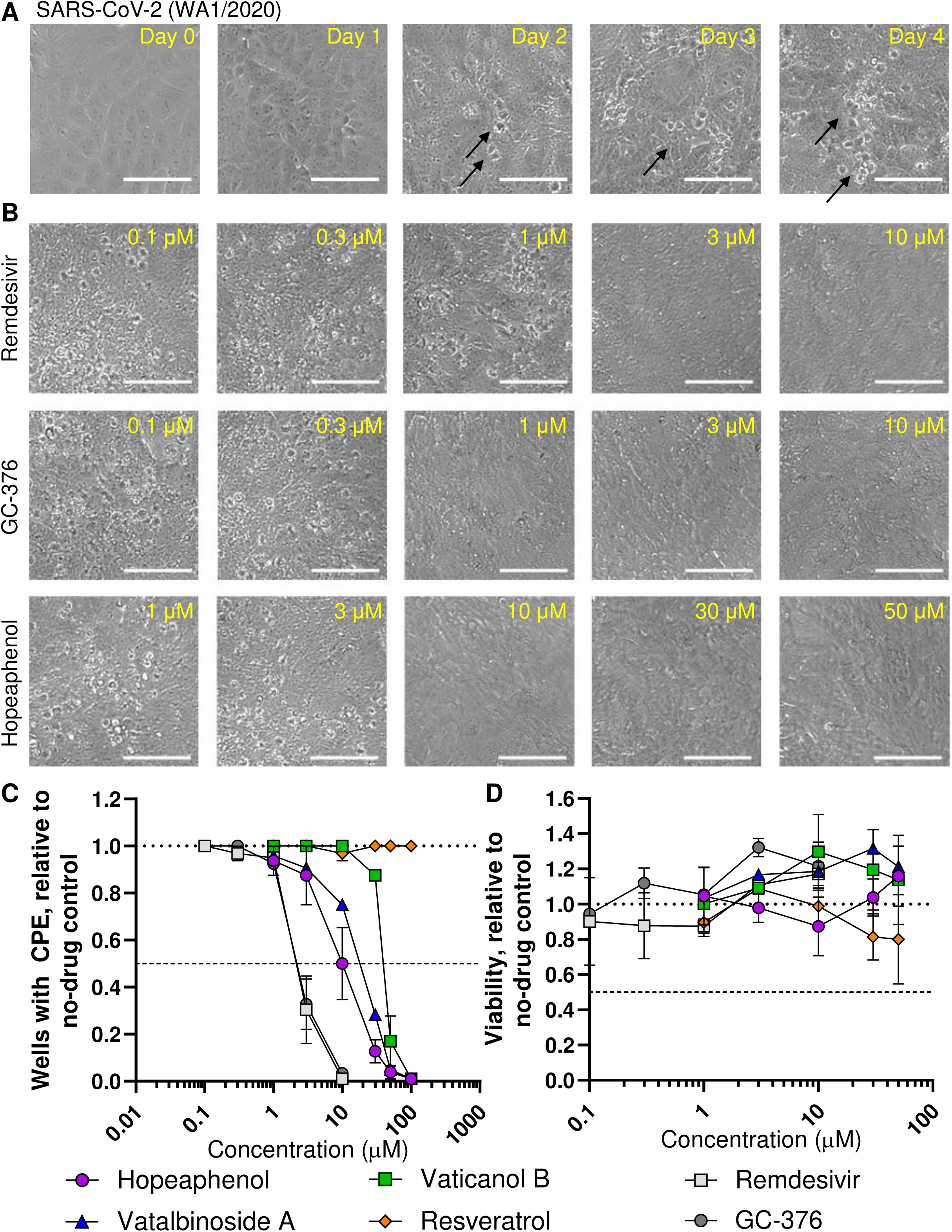
Effects of stilbenoids on infectious SARS-CoV-2 replication *in vitro*. **A,** Representative images of Vero-E6 cells infected with SARS-CoV-2 (USA-WA1/2020 variant) at 0 to 4 days post-infection. Arrows denote examples of CPE. **B,** Representative images of infected cells in the presence of remdesivir (top), GC-376 (middle), and hopeaphenol (bottom) after 4 days incubation at stated concentrations. Scale bars = 100 μm. **C,** Dose-response curves of stilbenoids, resveratrol, and remdesivir and GC-376 controls on viral replication in Vero-E6 cells after 4 days infection. **D,** Dose-response curves of compounds on cell viability in uninfected Vero-E6 after 4 days infection. In **C** and **D**, data are presented relative to cells treated with 0.1% DMSO vehicle control.

### (–)-Hopeaphenol inhibits SARS-CoV-2 variants of concern with improved efficacy against B.1.351

To determine if hopeaphenol maintained activity against emerging SARS-CoV-2 variants with accumulated mutations in the spike RBD, we repeated the CPE assay using two SARS-CoV-2 variants of concern including B.1.1.7 (England/204820464/2020; “UK variant”) and B.1.351 (KRISP-K005325/2020; “South Africa variant”) (**Figure 6**). Similar to our previous observations, infection of Vero-E6 cells with either strain resulted in widespread CPE across cultures after 4 days (**Figure 6A**, top), which was completely abolished by pre-treatment with 3 μM remdesivir (**Figure 6A**, middle). We also observed dose-dependent inhibition of these two variants by remdesivir, with calculated EC50s of 1.5 and 1.4 μM for B.1.1.7 and B.1.351 strains, respectively (**Figure 6B**), which also approximated observations with USA-WA1/2020 virus (EC50 = 2.5 μM; Table 1). Notably, 15 μM hopeaphenol also completely abrogated CPE by both variants (**Figure 6A**, bottom). When assessed for dose-response profiles, we observed that B.1.1.7 was inhibited by hopeaphenol with an EC50 of 14.8 μM (**Figure 6C**), which approximated hopeaphenol’s efficacy against USA-WA1/2020 (IC50 = 10.2 μM; **Table 1**). In contrast, B.1.351 was inhibited by hopeaphenol with an IC50 of 2.3 μM (**Figure 6C**), indicating 4.5-fold improved efficacy over USA-WA1/2020 (**Table 1**).

**Figure 6.**
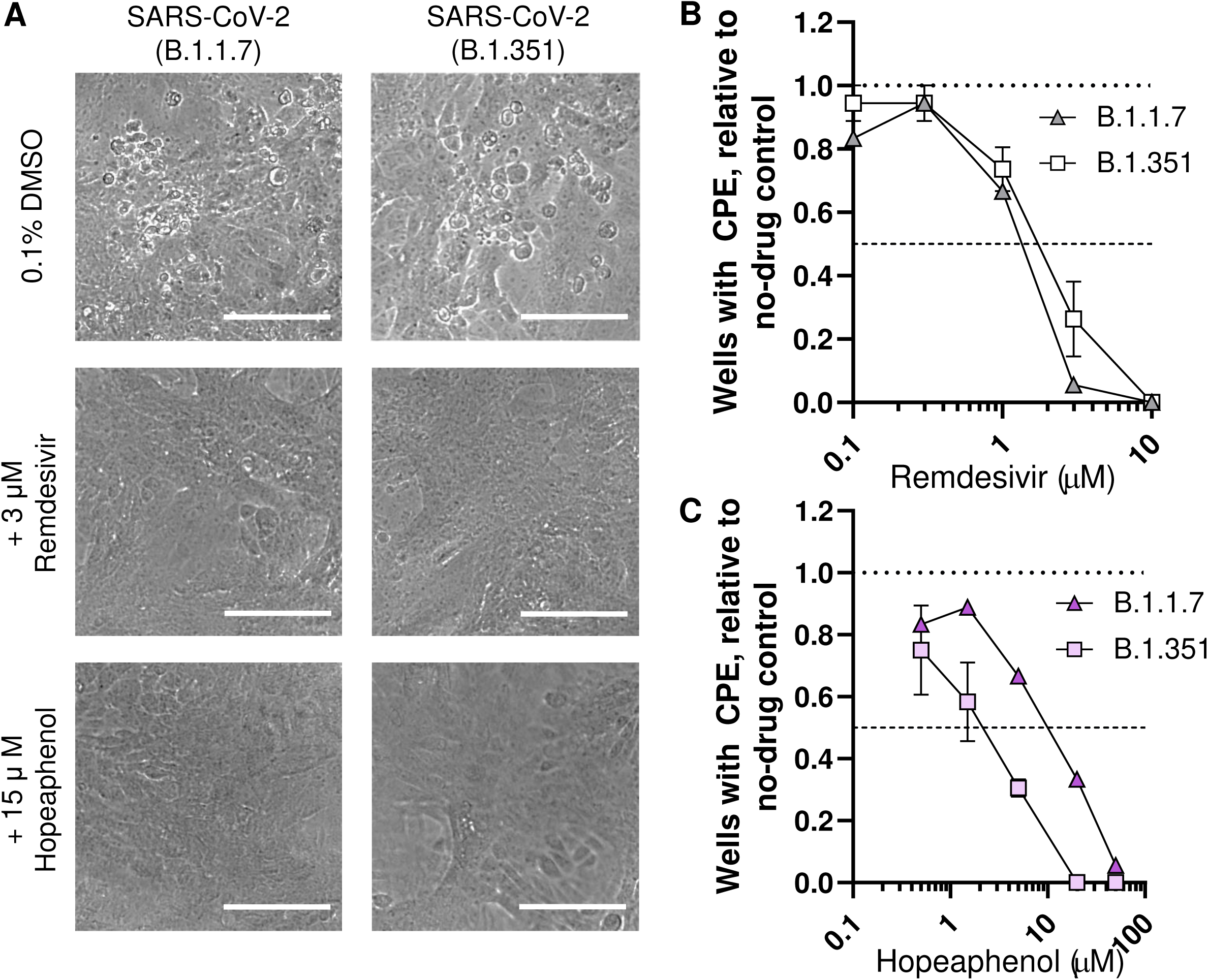
Effects of hopeaphenol and remdesivir on SARS-CoV-2 variant replication *in vitro*. **A,** Representative images of Vero-E6 cells following 4 days infection with either SARS-CoV-2 variants B.1.1.7 (left) or B.1.351 (right) in the presence of 0.1% DMSO vehicle control (top), 3 μM remdesivir (middle), or 15 μM hopeaphenol (bottom). Scale bar = 100 μm. **B-C,** Dose-response curves of remdesivir **(B)** or hopeaphenol **(C)** in Vero-E6 cells following 4 days infection with SARS-CoV-2 variant B.1.1.7 or B.1.351. In **B** and **C**, data are presented relative to infected cells treated with 0.1% DMSO vehicle control.

To confirm that the antiviral activities of hopephanol against these variants of concern corresponded to inhibition of viral entry, we generated VSVΔG-S-GFP-based pseudoviruses containing B.1.1.7 or B.1.351 spike sequences and infected Vero-E6 cells in the absence or presence of 50 μM hopeaphenol (**Figure 7**). Similar to previous observations (**Figure 4B**), no major changes in total cell nuclei number were observed under any experimental condition (data not shown). Also broadly consistent with previous observations with our original pseudovirus, we observed an average of 8.2 ± 2.9 and 6.0 ± 0.5% GFP-positive cells following 24 hours’ incubation with pseudovirus containing B.1.1.7 or B.1.351 spike, respectively (**Figure 7A**, top). Furthermore, infection of both pseudoviruses continued to be inhibited by 50 μM hopeaphenol (**Figure 7A**, bottom). For example, pseudovirus containing B.1.1.7 spike was observed in only 22.9 ± 7.6% of cells relative to infected, vehicle-treated cells treated with 0.1% DMSO (**Figure 7B**), similar to results from pseudovirus plus USA-WA1/2020 spike (28.4 ± 2.1%). In contrast, pseudovirus containing B.1.351 spike was present in only 15.3 ± 1.3% of infected, vehicle-treated cells when compared to infected cells without 50 μM hopeaphenol treatment (**Figure 7B**), indicating a 1.9-fold improved efficacy over USA-WA1/2020 spike-containing pseudovirus.

**Figure 7.**
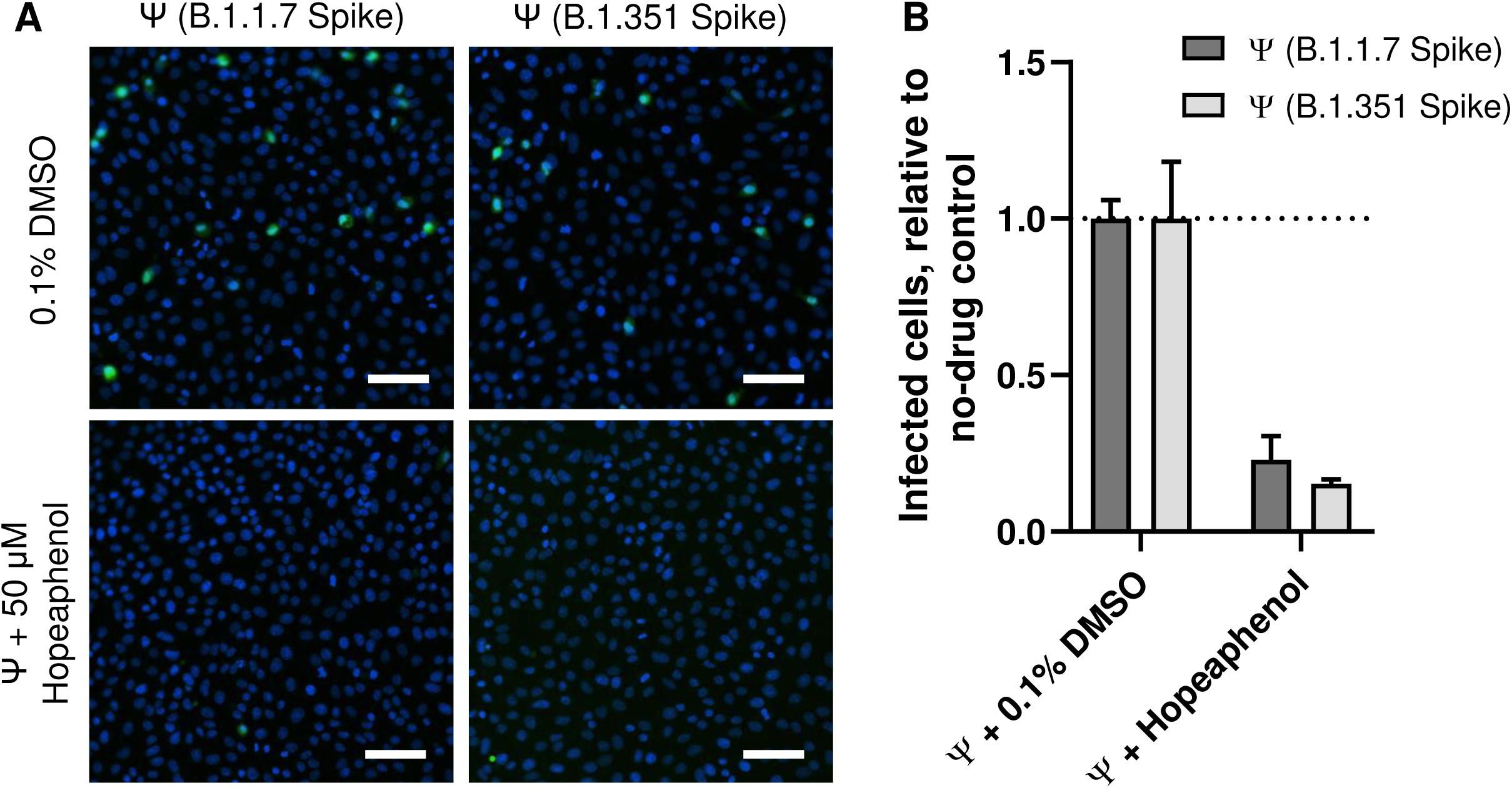
Effects of hopeaphenol on entry of pseudoviruses containing SARS-CoV-2 spike variants. **A,** Representative images of Vero-E6 cells infected with VSVΔG-S-GFP pseudovirus containing SARS-CoV-2 Spike from B.1.1.7 (left) or B.1.351 variants (right) in the presence of 0.1% DMSO (top) or 50 μM hopeaphenol (bottom). Images are organized as described in Figure 4. **B,** Level of GFP-positive (i.e., pseudovirus-infected) cells, relative to total cell nuclei, in the presence of 0.1% DMSO or 50 μM hopeaphenol.

Taken together, these results indicate that hopeaphenol inhibits both replication of infectious SARS-CoV-2 variants of concern *in vitro* and entry of pseudoviruses containing divergent SARS-CoV-2 spike sequences as well as improved efficacy against the B.1.351 variant.

## Discussion

Antivirals that act across multiple SARS-CoV-2 variants are needed worldwide to supplement emerging vaccine efforts. Here we investigated a library of pure compounds derived from 512 pure natural products and derivatives and identified three stilbenoids, exemplified by (–)-hopeaphenol, that disrupt the interaction of viral spike RBD with its host ACE2 receptor, block viral entry of spike-containing pseudoviruses, and antagonize infectious SARS-CoV-2 replication without cytotoxicity *in vitro*. Importantly, hopeaphenol also inhibits two emerging variants of concern (B.1.1.7 and B.1.351) which have acquired sequence variations that enhance SARS-CoV-2 infectivity and/or promote reduced susceptibility or escape from neutralizing antibodies. Hopeaphenol and other stilbenoid analogues are therefore promising leads for developing broad-spectrum SARS-CoV-2 entry inhibitors, potentially for use as monotherapies or in combination with antiviral leads against other viral targets.

(–)-Hopeaphenol, vatalbinoside A, vaticanol B, and related stilbenoids and their stereoisomers have been isolated from a variety of plant sources including *Hopea*, *Vitis*, *Shorea*, *Anisoptera*, and *Vatica* species, among others (29–37). These compounds have been reported to exhibit several *in vitro* properties including antiproliferative (32-33, 37-38), antibacterial (through inhibition of the type III secretion system of Gram-negative bacteria) (30, 39), antifungal (40), anti-influenza and herpes simplex virus (41–42), and anti-inflammatory (34, 43) activities, among others. Notably, hopeaphenol is additionally reported to inhibit plasma triglyceride elevation in olive oil-treated mice and reduce plasma glucose in sucrose-loaded mice at 200 mg/kg (44–45). It also exhibited hepatoprotective effects against LPS-induced liver injury in mice at 100 mg/kg (34). These initial *in vivo* efficacy studies, which indicate tolerability at high concentrations, support near-term *in vivo* studies of anti-SARS-CoV-2 efficacy by (–)- hopeaphenol.

More recently, stilbenoids have been proposed as potential disruptors of SARS-CoV-2 spike protein with ACE2 in molecular docking studies (46). Another stilbenoid, kobophenol A, was also recently reported to inhibit binding of RBD with ACE2 (IC50 = 1.8 μM) and SARS-CoV-2 replication (EC50 = 71.6 μM) (47), and our data are consistent with these observations. However, we observed no antiviral activity by resveratrol at up to 100 μM, which contrasts with another recent study reporting an EC50 of 10.7 μM in SARS-CoV-2-infected Vero-E6 cells (BetaCov isolate) (48). However, this latter study measured supernatant viral RNA levels by quantitative PCR after 48 hours’ infection, and so disparate results could reflect differences in sensitivity between quantitative PCR and CPE-based assays. Another consideration is that the three hit stilbenoid compounds, as polyphenolics, represent a structure class that has been given a PAINS (Pan Assay Interference compoundS) designation (49). Although we observed that these compounds selectively disrupted RBD/ACE2 binding over an unrelated PD-1/PD-L1 ligand/receptor pair, for example with 257.3-fold selectivity for hopeaphenol, caution must still be taken when considering these compounds for further therapeutic development. However, generation or isolation of stilbenoid analogues with potentially improved selectivity as well as assessment of chemical leads in primary cell models remain warranted.

While the three stilbenoids identified here selectively disrupted RBD/ACE2 interactions at sub-micromolar concentrations over an unrelated PD-1/PD-L1 ligand/receptor pair, and we observed efficacy at low micromolar concentrations in CPE assays, inhibitory activity against spike-containing pseudoviruses occurred only at 50 μM. These observations could partially reflect reduced stability of these stilbenoids *in vitro* and/or reduced efficacy against VSV- backbone pseudoviruses in particular. Stilbenoids are also sensitive to oxidation due to the presence of the phenolic moieties and their ability to delocalize an unpaired electron (50) They are also unstable to factors including oxygen, heat, light and pH changes (51–52). Consistent with the potential for reduced stability, vaticanol B, while consistently the most potent disruptor of RBD/ACE2 interactions by AlphaScreen, was ∼3-fold less effective than hopeaphenol and vatalbinoside A in both pseudovirus and CPE assays (**Table 1**). Assessment of additional analogues and derivatives may also mitigate this concern.

A recent report also describes use of a virtual screening approach which identified hopeaphenol as a potential inhibitor of SARS-CoV-2 M^pro^ by interacting within its active site (18). In contrast, we observed only weak inhibitory activity of hopeaphenol and analogues against M^pro^ (e.g., IC50s = ∼40 – 70 μM, compared to 0.0052 μM for GC-376; **Table 1**). While our studies do not rule out modest inhibition of M^pro^ by hopeaphenol, this activity is unlikely to confer the primary antiviral activity observed *in vitro* (e.g. hopeaphenol EC50s = 2.3 – 10.2 μM in CPE assays; **Table 1**). Support for this hypothesis also comes from our observations that hopeaphenol has similarly improved activity against the B.1.351 variant in both CPE and pseudovirus assays, indicating that hopeaphenol’s efficacy is dependent on spike sequence. Nevertheless, these combined results do raise the intriguing possibility of identifying stilbenoid derivatives that target both SARS-CoV-2 entry and M^pro^, which in turn may improve antiviral efficacy and/or reduce the risk of eventual viral drug resistance.

There are currently no licensed antivirals that reliably protect against COVID-19. Recent reports of SARS-CoV-2 variants with accumulated spike mutations and reduced susceptibility or escape from neutralizing sera from convalescent and vaccine-treated patients (7–8) also raise the concern of emerging variants with resistance to existing vaccines. In contrast, we observe that hopeaphenol, despite acting as an inhibitor of spike-mediated viral entry, inhibits CPE of both an early SARS-CoV-2 isolate (USA-WA1/2020) as well as two recently emerging variants of concern (B.1.1.7 and B.1.351), with improved efficacy against the antibody escape variant B.1.351. These results suggest that spike mutations that promote vaccine-induced viral escape may be distinct from those that might arise from ongoing treatment with hopeaphenol and potentially other stilbenoid-based entry inhibitors. Although further studies are clearly needed, this possibility, in turn, raises the possibility of natural product-based entry inhibitors that function as effective antiviral countermeasures in the absence of available second-generation vaccines.

## Materials and Methods

### Chemical libraries and hit compounds

The Davis Open Access Natural Product-Based Library consists of 512 distinct compounds, the majority (53%) of which are natural products obtained primarily from Australian fungal, plant, and marine invertebrate sources (11–12), as well as semi-synthetic natural product analogues (28%) and known commercial drugs or synthetic compounds inspired by natural products (19%). All compounds evaluated in this study were analyzed for purity prior to testing and shown to be > 95% pure. Compounds were initially provided by Compounds Australia at Griffith University in 5 mM stock solutions dissolved in dimethyl sulfoxide (DMSO; Millipore, Burlington, MA, USA); as such, DMSO was used as the vehicle control in this study. The three hit compounds identified following library screening, which are all known stilbenoids (30), were re-supplied as dry powders for confirmation studies and further biological evaluation.

### Cells, viruses, and reagents

Vero-E6 cells were obtained from the American Tissue Culture Collection. Vero-E6 cells were cultured in D10+ media [Dulbecco’s Modified Eagle Medium with 4.5 g/L glucose and L-glutamine (Gibco, Gaithersburg, MD), 10% fetal bovine serum (Gemini Bio Products, West Sacramento, CA, USA), 100 U of penicillin/mL, and 100 μg of streptomycin/mL (Sigma-Aldrich, St. Louis, MO)] in a humidified incubator at 37 °C and 5% CO_2_. BHK-21/WI-2 cells were purchased from Kerafast (Boston, MA, USA) and cultured in D5+ media, which is identical to D10+ media except for addition of 5% fetal bovine serum.

The following reagent was deposited by the Centers for Disease Control and Prevention and obtained through BEI Resources, NIAID, NIH: SARS-Related Coronavirus 2, Isolate USA-WA1/2020, NR-52281. The following reagents were obtained through BEI Resources, NIAID, NIH: SARS-Related Coronavirus 2, Isolate hCoV-19/England/204820464/2020, NR-54000, contributed by Bassam Hallis and SARS-Related Coronavirus 2, Isolate hCov-19/South Africa/KRISP-K005325/2020, NR-54009, contributed by Alex Sigal and Tulio de Oliveira.

Remdesivir and resveratrol were purchased from Sigma-Aldrich. GC-376 was purchased from Selleckchem (Houston, TX, USA). Isolation, structural confirmation, and purity of (-)- hopeaphenol, vatalbinoside A, and vaticanol B used in this study were reported previously (30).

### Protein-protein interaction assays

SARS-CoV-2 Spike-RBD binding to ACE2 was determined using AlphaScreen technology. 2 nM of ACE2-Fc (Sino Biological, Chesterbrook, PA, USA) was incubated with 5 nM HIS-tagged SARS-CoV-2 Spike-RBD (Sino Biological) in the presence of 5 μg/mL nickel chelate donor bead in a total of 10 μL of 20 mM Tris (pH 7.4), 150 mM KCl, and 0.05% CHAPS in white, opaque, low-volume 384-well plates. Test compounds were diluted to 100x final concentration in 100% DMSO. 5 μL of ACE2-Fc/Protein A acceptor bead was first added to the plate, followed by 100 nL of test compounds and then 5 μL of CoV-Spike-RBD-HIS/Nickel chelate donor beads. Test compounds were added to each well using a Janus Nanohead (PerkinElmer, Waltham, MA, USA). For each experiment, all conditions were performed in duplicate. Following 2 h incubation at room temperature, AlphaScreen fluorescent signals were measured using a ClarioStar plate reader (BMG Labtech, Cary, NC, USA). Data were normalized to percent inhibition, where 100% equaled the AlphaScreen signal in the absence of SARS-CoV-2-Spike-RBD-His and 0% equaled AlphaScreen signal in the presence of both proteins and DMSO alone.

PD-1 binding to PD-L1 was also determined using AlphaScreen technology. 0.5 nM of human PD-L1-Fc (Sino Biological) was incubated with 5 nM HIS-tagged human PD-1 (Sino Biological) in the presence of 5 μg/mL protein A AlphaScreen acceptor bead and 5 μg/mL nickel chelate donor bead in a total volume of 10 μL of 20 mM HEPES (pH 7.4), 150 mM NaCl, and 0.005% Tween in white, opaque low-volume 384-well plates. 5 μL of PD-L1-Fc/protein A acceptor bead was first added to the plate, followed by 100 nL of test compound prepared as described above, followed by 5 μL of PD-1-His/nickel chelate donor bead. For each experiment, all conditions were performed in duplicate. Following 2 h incubation at room temperature, data were collected as described above and normalized to percent inhibition, where 100% equaled the AlphaScreen signal in the absence of PD-1-His, and 0% equaled AlphaScreen signal in the presence of both proteins and 0.1% DMSO alone.

### Generation of M^pro^ protein

The codon-optimized gene for SARS-CoV-2 *M^pro^* (or *3CL^pro^*) (GenBank: QHD43415.1 aa 3264-3567) from strain BetaCoV/Wuhan/WIV04/2019 was ordered from IDT (Coralville, IA, USA) and cloned into a HIS-SUMO expression vector (a modified pET-DUET; Novagen, Madison, WI, USA). After transformation into BL21(DE3), the HIS-SUMO-M^pro^ fusion protein was expressed using the autoinduction method (53) with 500 mL cultures at 22 °C overnight. Cell pellets were resuspended in a buffer containing 25 mM Tris pH 8.5, 20 mM imidazole 200 mM NaCl, 5 mM b-mercaptoethanol and lysed using sonication and lysozyme and centrifuged at high speed. The supernatant was applied to a Ni-NTA (nickel-nitrilotriacetic acid) column at 4 °C and washed with the resuspension buffer. The fusion protein was then eluted using a buffer 300 mM imidazole, 200 mM NaCl and 5 mM b-mercaptoethanol, concentrated and applied to a gel filtration column (HiLoad 26/60 Superdex 75; Cytiva, Marlborough, MA) and equilibrated with the resuspension buffer. Fractions with > 90% purity were pooled and incubated with SUMO protease at 4 °C overnight. After cleavage, the digested protein solution was applied twice to a 5 mL HIS-TRAP Ni-NTA column (Cytiva) to remove the HIS-SUMO and SUMO protease, and the flow-through was collected. Finally, the protein was concentrated and applied to a second gel filtration column (HiLoad 26/60 Superdex 75; Cytiva) equilibrated with 25 mM HEPES pH 7.5, 150 mM NaCl, 2 mM TCEP. Purity (> 95%) was confirmed using an SDS-PAGE gel.

### M^pro^ enzymatic assays

Protease activity of recombinant M^pro^ was measured using the quenched fluorogenic substrate {DABCYL}-Lys-Thr-Ser-Ala-Val-Leu-Gln-Ser-Gly-Phe-Arg-Lys-Met-Glu- (EDANS)-NH2 (Bachem, Vista, CA, USA). 5 μL of 25 nM M^pro^ diluted in assay buffer [25 mM HEPES (pH 7.4), 150 mM NaCl, 5 mM DTT, 0.005% Tween) was dispensed into black, low-volume 384-well plates. Test compounds were serially diluted into 100% DMSO, and 0.1 μL was added to the assay using a Janus MDT Nanohead (PerkinElmer). Assays were initiated by addition of 5 μL of 5 μM fluorogenic substrate, and fluorescence at 355 nm excitation and 460 nm emission was monitored every 5 minutes for 50 minutes using an Envision plate reader (PerkinElmer). Rate of substrate cleavage was determined using linear regression of the raw data values obtained during the time course. Slopes of these progress curves were then normalized to percent inhibition, where 100% equaled rate in the absence of M^pro^ (which was typically 0), and 0% equaled rate of cleavage in the presence of M^pro^ and 0.1% DMSO.

### Generation of VSVΔG-S-GFP pseudoviruses

SARS-CoV-2 USA-WA1/2020 cDNA was obtained as a gift from Dr. Stephen J. Elledge. B.1.1.7 spike cDNA was generated from USA-WA1/2020 spike cDNA by standard PCR mutagenesis, and B.1.351 spike cDNA was synthesized (Genscript, Piscataway, NJ, USA). Pseudoviruses were generated in BHK-21/WI-2 cells using a pseudotyped ΔG-GFP (G*ΔG-GFP) rVSV (Kerafast, Boston, MA, USA) in addition to spike sequences cloned into the paT7- Spike plasmid as described previously (21). 2 hours following chemical transfection of spike plasmid, cells were infected with ΔG-GFP (G*ΔG-GFP) rVSV. Following 24 hours incubation, supernatants were harvested, aliquoted, and stored at -80 °C.

### Pseudovirus-based infectivity assays

12,500 Vero-E6 cells resuspended 12.5 μL D10+ were plated in 384 μL plates, followed by addition of 6.25 μL of test agents diluted in D10+ at 4X desired final concentration plus 6.25 μL of undiluted pseudovirus stock (total 25 μL reaction volumes). All experimental conditions with test agents were performed in duplicate, and control cells with or without pseudovirus in the presence of 0.1% DMSO vehicle control were tested 4-fold. Cells were incubated at 37 °C and 5% CO_2_ for 24 hours. Cells were then stained with 25 μL of 5 μg/mL Hoechst 33342 (Sigma Aldrich), incubated for 20 minutes, and fixed with paraformaldehyde to 2% final concentration. High content imaging was then performed using a Nikon Eclipse Ti Inverted Microscope and Nikon NIS Elements AR Software Version 5.30.02 (Nikon Americas Inc., Melville, NY, USA). For each image, cell nuclei and GFP-positive cells were counted, with GFP positive cells reported as percent of total nuclei within each image.

### Resazurin cell viability assay

20,000 Vero-E6 cells were plated in 96-well plates and incubated overnight before addition of compounds at defined concentrations. 0.1% DMSO vehicle control was added to wells in the absence of test compounds. All experimental conditions were performed in duplicate. Cells were then incubated at 37 °C and 5% CO_2_ for 4 days before addition of resazurin (Sigma Aldrich) to a final concentration of 20 μg/mL. Cells were incubated for an additional 4 hours before fluorescence intensity was measured using a ClarioStar plate reader (BMG Labtech). Background fluorescence was subtracted from wells containing resazurin and D10+ media but no cells.

### Generation of SARS-CoV-2 viruses

3 x 10^6^ Vero-E6 cells were incubated in 15 mL of D10+ media for 24 hours. Cells were then washed and replaced with 10 mL of D10+ containing virus at a multiplicity of infection MOI of 0.001. Cells were incubated for 5 – 7 days until clear CPE was observed throughout the flask. Media was harvested and split into 250 μL aliquots for storage at -80 °C.

To titer virus stocks, Vero-E6 cells were first plated to 20,000 cells per well in 96-well format in D10+ media and incubated for 24 hours. Following incubation, cells were incubated in fresh D10+ containing 5-fold serial dilutions of a thawed virus aliquot (8 total dilutions, 5-fold replicates) and incubated for an additional 4 days. Wells were then scored for the presence of CPE. TCID50s were calculated using the Reed-Muench method.

### Viral CPE assays

Vero-E6 cells were plated in D10+ to 20,000 cells per well in 96-well format and incubated for 24 hours. Following incubation, compounds were added to final concentrations in 8-fold replicates and incubated for an additional 2 hours before addition of 50x TCID50 of virus. Each 96-well plate further contained uninfected cells and infected cells with 0.1% DMSO vehicle control in 4-fold replicates. Cells were incubated for an additional 4 days, at which point all wells were scored for the presence of CPE by a user blinded to the identity of the wells.

### Data analysis

For all studies, 50% effective concentrations were calculated using nonlinear regression of a one-side binding model using GraphPad Prism v. 8.4.3 (GraphPad, San Diego, CA, USA). All data are presented as the mean ± s.e.m. from at least 3 independent experiments.

## Acknowledgements

We are indebted to Dr. Stephen J. Elledge for providing SARS-CoV-2 spike cDNA for this study. Funding was provided by the Wistar Science Discovery Fund (L.J.M., J.S.), Canadian Institutes for Health Research (CIHR PJT-153057) (I.T.) and a Griffith University–Simon Fraser University Collaborative Travel Grant (I.T., R.A.D.). The authors acknowledge the National Health and Medical Research Council (APP1024314 to R.A.D.), and the Australian Research Council for support towards NMR and MS equipment (LE0668477, LE140100119, and LE0237908) and a linkage research grant (LP120200339 to R.A.D.). R.A.D. acknowledges the NatureBank biota repository (https://www.griffith.edu.au/institute-drug-discovery/unique-resources/naturebank) from which the majority of compounds found within the Davis open access natural product-based library were purified. Compounds Australia (https://www.griffith.edu.au/griffith-sciences/compounds-australia) is acknowledged for curating, plating and shipping the Davis open access library. This work was also supported by the following grants to L.J.M.: the Robert I. Jacobs Fund of The Philadelphia Foundation and the Herbert Kean, M.D., Family Professorship.

